# An Optimal Lysis Time Maximizes Bacteriophage Fitness in Quasi-continuous Culture

**DOI:** 10.1101/2020.05.11.089508

**Authors:** Sherin Kannoly, Abhyudai Singh, John J. Dennehy

## Abstract

Optimality models have a checkered history in evolutionary biology. While optimality models have been successful in providing valuable insight into the evolution of a wide variety of biological traits, a common objection is that optimality models are overly simplistic and ignore organismal genetics. We revisit evolutionary optimization in the context of a major bacteriophage life history trait, lysis time. Lysis time refers to the period spanning phage infection of a host cell and its lysis, whereupon phage progeny are released. Lysis time, therefore, directly determines phage fecundity assuming progeny assembly rate is maximized. Noting that previous tests of lysis time optimality rely on batch culture, we implemented a quasi-continuous culture system to observe productivity of a panel of isogenic phage λ mutants differing in lysis time. We report that λ phage productivity in our experiments is maximized around an optimal lysis time of 63 min, which is the lysis time of the λ “wildtype” strain. We discuss this finding in light of recent results that lysis time variation is also minimized in the λ “wildtype” strain.

## Introduction

Evolutionary biologists have long been interested in the power of natural selection to refine biological adaptations (Smith 1978; Endler 1986; Lewontin 1989; Crow 1993). Generally, views on the power of natural selection fall into two camps: one emphasizing the ability of natural selection to optimize biological traits (Endler 1986; Parker and Smith 1990) and the other stressing the limitations of natural selection in face of evolutionary tradeoffs and genetic details (Lewontin 1989; Crow 1993) affecting phenotypic traits. Several studies have employed bacteriophages applied optimization principles to theoretically predict how these traits will evolve in response to changing ecological conditions (Abedon 1989; Wang et al. 1996; Abedon et al. 2001; Bull et al. 2004; Bull 2006; Bonachela and Levin 2014).

However, despite clear predictions, experimental tests have generally failed to confirm theoretical predictions. T7, ST-1, and ΦX174 phages with theoretically predicted suboptimal lysis times generally failed to evolve predicted optimal lysis times (Heineman et al. 2007; Chantranupong and Heineman 2012). While the evolution of T7 phage with a deleted lysin gene qualitatively supported optimality predictions, it was noted that abolition of the lysis function did not extend the latent period, thus had no effect on the expected tradeoff between latent period and burst size (Heineman et al. 2007). ST-1 qualitatively, but not quantitatively, adapted as predicted by modeling, and ΦX174 failed to adapt at all (Chantranupong and Heineman 2012). In another study, experimental evolution of phage RB69 did result in selection for shorter lysis time mutants in high host density cultures, but it is unclear whether the lysis time of these mutants were optimal for the experimental conditions (Abedon et al. 2003). A study employing isogenic lambda phages with different lysis times found the model predictions for both the optimum and fitness to be different from the experimental estimations (Wang 2006).

However, we note that these studies used batch cultures to examine host-phage interactions, and phage fitnesses were estimated using absolute growth rates. These conditions may not effectively mimic the kinds of conditions experienced by phages in their natural habitats. To revisit the question of phage optimal lysis time, we employed a different approach, which was inspired by Bonachela *et. al* (2014), who modeled host-virus interactions in natural environments in terms of steady state conditions, such as those seen in continuous culture systems. A typical example of a continuous culture system is a two-stage chemostat. In the first stage, a chemostat receives a flow of fresh media to maintain a continuous culture of bacteria. Fresh host bacteria from the first chemostat are fed into a second chemostat containing a population of phages. Both host cells and phages are washed out from the second chemostat at a specified rate. This system can mimic steady states that exist in natural environments, such as the gastrointestinal tract where peristaltic movements can maintain a continuous flow.

In this study, we maintained phage λ and its host, *Escherichia coli*, under quasi-steady state conditions. To initiate experiments, λ holin mutants differing in their lysis times (Dennehy and Wang 2011; Singh and Dennehy 2014; Kannoly et al. 2020) were added to exponentially replicating batch cultures of *E. coli*. After a short period of growth, λ phages were filtered from host cells, and a small fraction was transferred into a fresh exponential culture. At no time did these phages experience stationary phase host cells during the course of the experiment. In principle, differences in lysis times should result in differences in phage productivity (Levin et al. 1977; Wang 2006). We predicted that there exists an optimum for lysis time that maximizes phage production under these quasi-steady state conditions. By contrast, strains with suboptimal lysis times should show declining phage populations in this experimental context. Our results show that phage production in these quasi-steady state conditions is maximized at lysis times approaching 63 min, which, perhaps not coincidentally, is the lysis time of the phage λ wildtype. These results suggest an optimum lysis time that maximizes phage progeny production.

## RESULTS AND DISCUSSION

Lysis is the last step in the infectious cycle of lytic bacteriophage, resulting in the release of virions. Lysis timing in many phage is controlled by a single protein, holin, which accumulates in the cell membrane up to a genetically determined time, whence it nucleates generating membrane-permeabilizing holes (Wang et al. 2000, 2003; Dennehy and Wang 2011; Singh and Dennehy 2014; Kannoly et al. 2020).

To study the effect of lysis time on phage productivity (i.e., total phage reproduction during a fixed growth period), we employed a panel of λ phages harboring mutations in the holin gene *S*, which delay or hasten the lysis event following induction of the lytic cycle. These λ mutants were used to initiate batch cultures containing a fixed concentration of *E. coli* cells at a multiplicity of infection (MOI) of 0.001, then purified, diluted 1000-fold, and transferred to fresh cultures at 3 h intervals, generating quasi-steady state conditions. A 3 h infection allows at least three generations for wild type λ such that the MOI ≈ 1 at the end of phage growth. A low MOI also minimizes multiple infections per cell and ensures multiple generations without depleting host cells to such an extent that the cell number becomes a limiting factor for phage growth. Furthermore, the short duration of growth prevents *E. coli* densities from being sufficiently high enough to enter stationary phase.

Ideally, this study would be performed using actual chemostats, but we choose not to pursue this option because it would limit the number of holin mutants and degree of replication that we could easily manage. Moreover, it is well known that *E. coli* growth is consistent (i.e., in a steady state) as long as exponential growth conditions are maintained (Fishov et al. 1995) so, in physiological terms, *E. coli* growth is not inherently different between chemostats and our quasi-steady state transfers. Such a steady state achieved by introducing fresh batch cultures ensures a balanced exponential phase throughout the transfers. The cells in exponential phase are in the same physiological state characterized by relatively similar cell volumes and macromolecular composition. This aspect ensures that the growth parameters such as cell density (number of cells per milliliter) and the concentration of key macromolecules such as proteins, RNA, DNA, etc., increase exponentially with precisely the same doubling time (Sezonov et al. 2007).

Pilot studies were performed by initiating wild type λ infections of host cells in exponential phase. Cells exceeded phage by 1000-fold minimizing the time required for phage to encounter host cells for attachment and ensuring that the MOI was close to one at the end of the first 3-h transfer. Experiments where the growth period was less than 3 h per transfer and the dilution between transfers was than 1000-fold resulted in lower phage numbers and increased likelihood of phage population extinction after a few transfer cycles, especially in fast lysing strains (data not shown). On the other hand, experiments where the growth period was longer than 3 h resulted in higher phage yields, and the host populations were depleted before the end of the growth period, especially in strains with long lysis times.

For the subsequent serial transfers after the initial growth period, phages were diluted 1000-fold and this dilution rate was kept constant throughout the transfers. The transfer conditions were adjusted using the wild type strain to obtain increasing productivity after each transfer until it peaks after the fifth transfer. Subsequent transfers did not result in any significant increase in productivity primarily due to coinfections and faster host cell depletion (data not shown). Moreover, five transfers would result in approximately fifteen phage generations, beyond which phage adaptation would begin to impact our results. It is important to note that unlike previous studies, our study does not aim to explore evolution of an optimum lysis time as a result of phenotypic adaptation. Instead, we aim to test the productivity (i.e., total phage reproduction during a fixed growth period) of genotypes varying in their lysis times under dynamic environments, which are subject to constant dilution or wash out. To this end, the experimental protocol was designed to mimic conditions were both phages and cells would be washed out at regular intervals.

Five such serial transfers were completed, and the phage titers were obtained after every transfer (Fig. 1). Fig. 2 shows phage titers after every transfer for all the strains and Fig. 3 compares phage productivity only for the fifth transfer for all strains. Our data show that phage progeny production following five transfers was maximized across a broad range spanning 60–100 min (Fig. 2; Fig. 3). Interestingly the phage λ wildtype strain falls within this range (Fig. 2; Fig. 3). Outside this range, phage progeny production drops precipitously up to six orders of magnitude (Fig. 2; Fig. 3). This outcome demonstrates the strong influence of lysis timing on the reproductive success of the phage.

**Fig 1.**
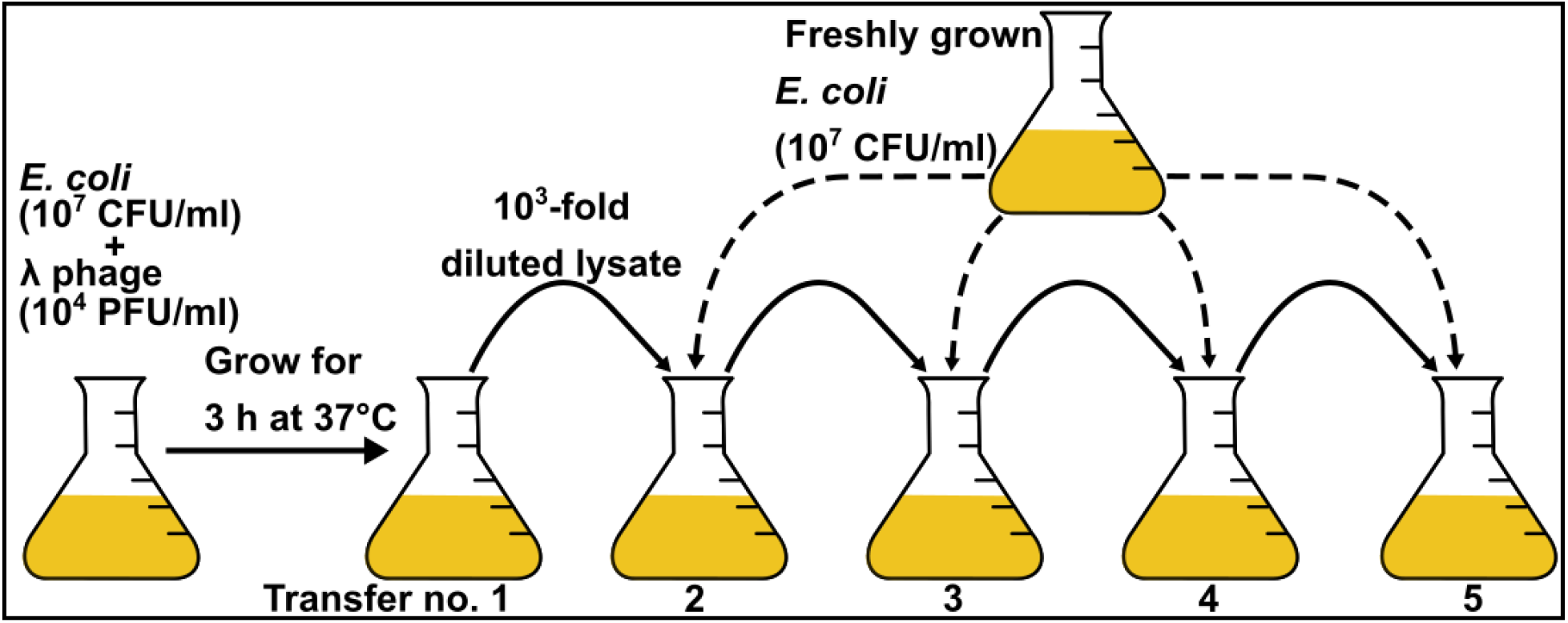
Serial transfers. For serial transfers, λ phage strains were cocultured with exponentially growing host cells, filtered to separate phages, and diluted 1000-fold to start the next transfer with freshly growing cultures. After every transfer, the filtered phage lysates were titered using plaque assays.

**Fig 2.**
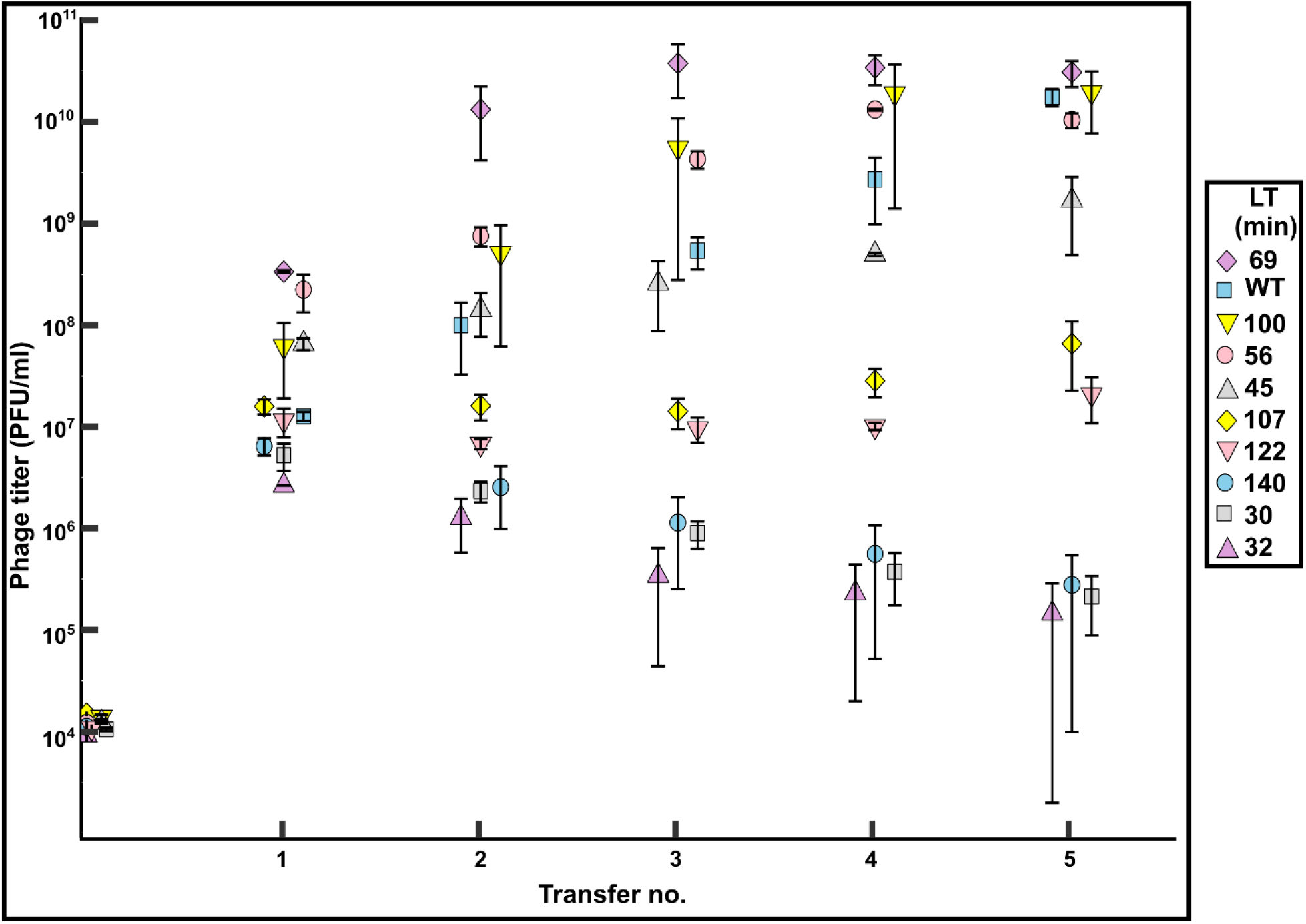
Titers of λ strains with different lysis times (LT) after each transfer. Lysis times are from single-cell estimates as reported in Kannoly et al., 2020. WT lysis time = 63 min. Error bars, mean ± SEM.

**Fig 3.**
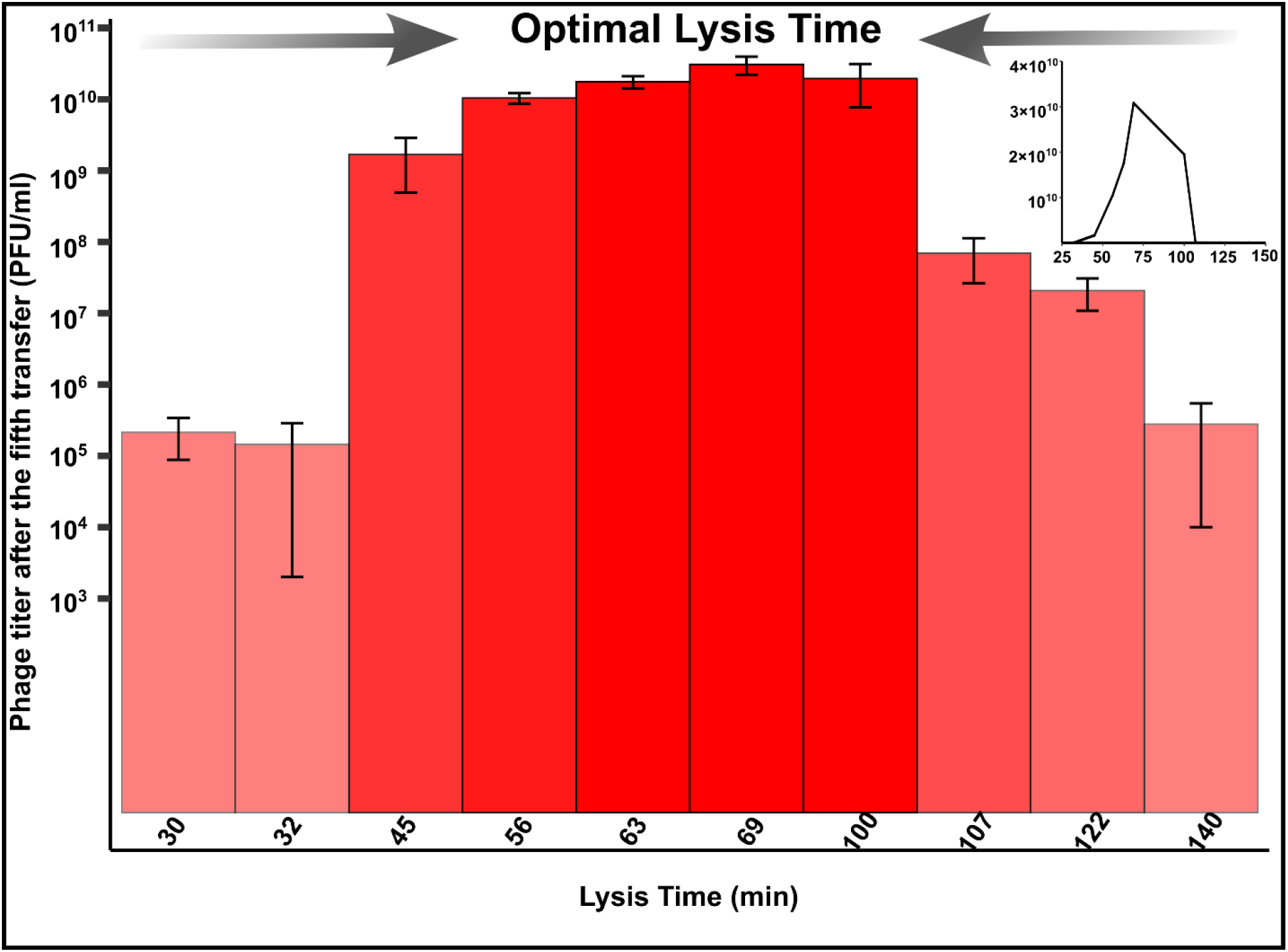
Titers of λ strains with different lysis times after the fifth transfer (Log scale). Titers increase as the lysis time approaches optima. Inset shows the same data on a linear scale. Error bars, mean ± SEM.

Conceptually, phage production per generation depends on the burst size, which is positively correlated to the lysis time (Wang 2006; Baker et al. 2016). The exact shape of this correlation is a matter of debate (Baker et al. 2016). A phage genotype with a short lysis time and hence a low burst size would result in an overall reduction in phage productivity. Thus, fewer phages are carried over to the next transfer. Conversely, one might intuitively assume that for delayed-lysing strains, the phage productivity will be significantly higher. However, the delay in lysing traps a significant number of phages within cells, which fail to make it to the next transfer. Although delaying lysis increases the number of progenies, it also delays the generation time. This trade-off makes the optimum sensitive to environmental conditions such as dilution due to a constant flow rate. Our experimental set up simulates such a dilution, and the strains with lysis times that deviate from the optimum show reduced phage productivity over subsequent transfers (Figs. 2 and 3). In a previous study using phage λ holin mutants, we demonstrated that holin accumulation thresholds generate precision in lysis timing. We showed that noise in lysis timing is reduced in mutants with lysis times closer to that of the wild type (Ghusinga et al. 2017; Kannoly et al. 2020). Current results support the idea of an optimal lysis time presumably derived from minimizing the variation in lysis timing. This is in stark contrast to previous studies, which have suggested that mutations that increase lysis time variance would be favored by selection (Bull et al. 2011; Baker et al. 2016). It was hypothesized that since phage burst size increase linearly (Hutchison and Sinsheimer 1966; Wang 2006), early lysis events would contribute more to population growth than late lysis events detract (Bull et al. 2011; Baker et al. 2016). Interestingly, Storms *et. al* (2014) have reported that T4 phage productivity in a cell about to undergo cell division was almost three times greater than the productivity in a young, newly formed cell. We speculate that if the contribution of early lysis events to population growth is balanced by larger bursts in the later stages of infection, mutations that increase lysis time variance may not be favored after all (Baker et al. 2016).

Taken together, minimizing lysis time variance may factor in the evolution of holin. In this study, we used holin mutants to further explore the effect of lysis time on phage productivity under quasi-steady state conditions. Again, mutants with lysis times closer to that of the wildtype showed a consistent increase in phage productivity compared to those with suboptimal lysis times. Optimality models can reveal the underlying ecological contexts influencing evolutionary processes. Our results suggest that the wildtype λ might already be selected for conditions where a constant dilution results in a continuous wash out. Further experiments to test this hypothesis might involve direct competition of wildtype against mutants under similar conditions. Also, long-term experimental evolution under the same conditions should reveal if phage strains with suboptimal lysis times could evolve towards optimality.

Our results add additional support for the idea that life history traits evolve to optimal values. Much work has been conducted experimentally assessing the validity of optimality models with varying degrees of success (Kozlowski 1999; Dekel and Alon 2005; Miller et al. 2006; Camps et al. 2007; Poelwijk et al. 2011b, 2011a; Nakabayashi 2012; Graves and Weinreich 2017). At this point, it is clear that an unnuanced account of optimality evolution may fail because of genetic details or constraints (Bull et al. 2004). Nonetheless, optimality models have provided valuable insights into biological traits ranging from the genetic code to animal behavior (Charnov 1976; Parker and Smith 1990; Jensen et al. 2006; Roff et al. 2006; Noblin et al. 2008; Baranov et al. 2009; Henshaw 2018; Ruess et al. 2019).

## MATERIALS AND METHODS

### Strains

All bacteria and lysogen strains used in this study are listed in Table 1.

**Table 1.** All bacteria and lysogen strains used in this study are listed along with the genotype of the prophage’s holin gene and the source of the strain. ^a^Coli Genetic Stock Center.

| Strain | Genotype | Lysis Time (min) | Source |
| --- | --- | --- | --- |
| CGSC#: 6152<sup>a</sup> | *E. coli* MC4100 (λ-) | NA | Casadaban 1976 |
| JJD3 | MC4100 ($\lambda$ *cI857 S*) | 63 | Wang 2006 |
| JJD246 | MC4100 ($\lambda$ *cI857 S105<sub>H7D</sub>*) | 69 | Kannoly et al., 2020 |
| JJD248 | MC4100 ($\lambda$ *cI857 S105<sub>F94C</sub>*) | 56 | Kannoly et al., 2020 |
| JJD391 | MC4100 ($\lambda$ *cI857 S105<sub>A16G/K92Q</sub>*) | 140 | Kannoly et al., 2020 |
| JJD404 | MC4100 ($\lambda$ *cI857 S105<sub>I21V</sub>*) | 30 | Kannoly et al., 2020 |
| JJD405 | MC4100 ($\lambda$ *cI857 S105<sub>V45G</sub>*) | 32 | Kannoly et al., 2020 |
| JJD414 | MC4100 ($\lambda$ *cI857 S105<sub>L90I</sub>*) | 45 | Kannoly et al., 2020 |
| JJD423 | MC4100 ($\lambda$ *cI857 S105<sub>F27Y</sub>*) | 122 | Kannoly et al., 2020 |
| JJD432 | MC4100 ($\lambda$ *cI857 S105<sub>S89W</sub>*) | 107 | Kannoly et al., 2020 |
| JJD436 | MC4100 ($\lambda$ *cI857 S105<sub>K92N</sub>*) | 100 | Kannoly et al., 2020 |

### Plaque assays

The plaque assays were designed for minimizing variation in plaque size and enabling plaque counting. The *E. coli* strain MC4100 was grown overnight in TB broth (5 g NaCl and 10 g tryptone in 1 L water) plus 0.2% maltose at 37°C. The overnight culture was diluted with equal volume of TB + maltose and grown for another 1.5 h. 100 ul of these cells were mixed with appropriately diluted phage lysates and incubated at room temperature for 20 min to allow pre-adsorption. This mixture was then added to 2.5 ml of molten H-top agar (Miller 1992), gently vortexed, and overlaid onto freshly prepared plates containing 35 ml LB agar. The plates were then incubated at 37°C and plaques were counted after 18–22 h. Each assay was triplicated to estimate the average plaque forming units per milliliters (PFU/ml).

### Thermal induction of lysogens to obtain phage lysates

Lysogens were grown overnight in LB media at the permissive temperature of 30°C. Overnight cultures were diluted 100-fold and grown in a 30°C shaking incubator till an OD_600_ of 0.3–0.4 was reached. For heat induction, the cultures were transferred to a 42°C shaking water bath for 20 min. Following induction, the cultures were transferred to a 37°C shaking incubator until lysis. The phage lysates were filtered and titered using plaque assay.

### Single-cell lysis time determination

We induced the lytic cycle in lysogens and recorded the lysis times of single cells to estimate the mean lysis time. The protocol for determining single-cell lysis times has been described previously (Dennehy and Wang 2011). Briefly, a 200-µl aliquot of exponentially growing culture (OD_600_ = 0.3–0.4) of lysogens was chemically fixed to a 22 mm square glass coverslip, which was pretreated with 0.01% poly-L-lysine (mol. wt. 150 K–300 K; Millipore Sigma, St. Louis, MO) at room temperature for 10 min. Using this coverslip, a perfusion chamber (RC-21B, Warner Instruments, New Haven, CT) was assembled and immediately placed on a heated platform (PM2; Warner Instruments, New Haven, CT). The heated platform was mounted on an inverted microscope stage (TS100, Nikon, Melville, NY), and the perfusion chamber was infused with heated LB at 30°C (Inline heater: SH-27B, dual channel heating controller: TC-344B; Warner Instruments, New Haven, CT). The chamber temperature was elevated to 42°C for 20 min and thereafter maintained at 37°C until ∼95% cells were lysed. Videos of cell lysis were recorded using an eyepiece camera (10X MiniVID^™^; LW Scientific, Norcross, GA). The lysis times of individual cells were visually determined using VLC™ media player. Lysis time was defined as the time required for a cell to disappear after the temperature was increased to 42°C. For each lysogen, an average of approximately 100 cells was used to calculate the lysis time.

### Serial Transfers

To initiate the transfers, an *E. coli* (∼10^7^ CFU/ml) culture growing exponentially at 37°C in a shaking incubator (200 rpm) was infected with a phage lysate (∼10^4^ PFU/ml). The infection could proceed for 3 h, after which phages were separated by filtration using a 0.2 µ syringe filter (Pall Corp.). The filtered lysate was diluted 1000-fold to start the next transfer using freshly growing cells (∼10^7^ CFU/ml). These steps were repeated four more times for a total of five serial transfers (Fig. 1). Plaque assays were performed at the end of each transfer to determine the titers of phage lysates. The transfers for all strains were duplicated.

## Acknowledgements

This work was funded by National Institutes of Health NIGMS grant 1R01GM124446-01 to AS and JJD. We thank Ry Young for stimulating conversations about phage λ, and for the phage and bacterial strains described herein. We also appreciate thought-provoking discussions about phage life history with Stephen Abedon, Jim Bull, Khem Ghusinga, Cesar Vargas-Garcia, Ing-Nang Wang, and Daniel Weinreich.

